# Highly regenerative species-specific genes improve age-associated features in the adult *Drosophila* midgut

**DOI:** 10.1101/2023.07.04.547653

**Authors:** Hiroki Nagai, Yuya Adachi, Tenki Nakasugi, Ema Takigawa, Junichiro Ui, Takashi Makino, Masayuki Miura, Yu-ichiro Nakajima

**Affiliations:** Graduate School of Pharmaceutical Sciences, The University of Tokyo; Graduate School of Life Science, Tohoku University

**Keywords:** Synthetic biology, aging, healthy lifespan, regeneration, *Drosophila*, intestinal stem cells

## Abstract

**Background:** The remarkable regenerative abilities observed in planarians and cnidarians are closely linked to the active proliferation of adult stem cells and the precise differentiation of their progeny, both of which typically deteriorate during aging in low regenerative animals. While regeneration-specific genes conserved in highly regenerative organisms may confer regenerative abilities and long-term maintenance of tissue homeostasis, it remains unclear whether introducing these regenerative genes into low regenerative animals can improve their regeneration and aging processes.

**Results:** Here we ectopically express high regenerative species-specific JmjC domain-encoding genes (HRJDs) in *Drosophila*, a widely used low regenerative model organism. Surprisingly, HRJD expression impedes tissue regeneration in the developing wing disc but extends organismal lifespan when expressed in the intestinal stem cell lineages of the adult midgut under non-regenerative conditions. Notably, HRJDs enhance the proliferative activity of intestinal stem cells while maintaining their differentiation fidelity, ameliorating age-related decline in gut barrier functions.

**Conclusions:** These findings together suggest that the introduction of highly regenerative species-specific genes can improve stem cell functions and promote a healthy lifespan when expressed in aging animals.

## INTRODUCTION

Regeneration, an intricate process that rebuilds lost body parts, is a widespread phenomenon among metazoans, but the capacity for regeneration displays significant variation across different groups and species [1–5]. While certain animals like planarians and hydras possess the remarkable ability to regenerate their entire body from a small fragment, other groups with more complex body structures, such as mammals and insects, exhibit a diminished regenerative potential and can only regenerate specific tissues and/or organs to a limited extent. Furthermore, regenerative capacity often declines with aging in most species with limited regeneration abilities [2], resulting in increased susceptibility to organismal death upon injury. In contrast, animals that can achieve whole body regeneration, along with developmental reversion observed in the jellyfish *Turritopsis*, exhibit potential immortality [2, 5, 6]. Understanding the mechanisms underlying high regenerative ability and their relationship with aging represents a fundamental challenge in the field of developmental biology and gerontology with implications for regenerative medicine.

Several cellular and molecular factors have been identified as determinants of regeneration capacity. Highly regenerative animals such as planarians and cnidarian polyps rely on pluripotent adult stem cells, called neoblasts and interstitial cells (i-cells), respectively [2–5, 7]. These stem cells migrate to the injury sites and contribute to the formation of a blastema, an undifferentiated cellular mass, enabling the restoration of amputated body structures. Some vertebrates like salamanders and fish, which do not possess adult pluripotent stem cells, can regenerate organs after injury by recruiting blastema cells through dedifferentiation and/or the activation of quiescent lineage-restricted stem cells [1, 2, 4, 5, 8]. At the molecular level, the evolutionary conserved WNT signaling pathway promotes a wide range of regenerative events across species, including blastema formation in newts and *Hydra* [1–5, 8].

In contrast to the conserved regulators of regeneration, several genes are specific to highly regenerative animal groups and species: for instance, the newt gene *Prod1* regulates re-patterning during limb regeneration [9, 10], and viropana family (*viropana 1-5*) is up-regulated during lens regeneration [11, 12]. These species/group-specific genes might explain differences in regeneration capacity between species. Remarkably, ectopic expression of *viropana 1-5* can enhance regeneration of the primordium of *Drosophila* eyes that maintain regenerative capacity during development [12]. This finding raises the possibility that heterologous induction of regenerative genes may accelerate tissue regeneration, at least in developing animals, and potentially provide a cue for developing novel regenerative therapies. However, it remains unknown whether heterologously-induced regenerative genes can improve regenerative and/or aging processes even when induced in post-developmental mature adults.

Notably, given that basal metazoans such as Porifera, Ctenophore, Placozoa, and Cnidaria all exhibit robust regenerative abilities, it is conceivable that a common ancestor of all metazoans once possessed a high regenerative potential and independently lost genes related to high regenerative capacity in multiple phyla. Building upon this hypothesis, bioinformatics analysis has identified genes that are common among species with high regenerative abilities and absent in species with limited regenerative capacities (Fig. 1A) [13]. The <u>h</u>ighly <u>r</u>egenerative species-specific <u>J</u>mjC <u>d</u>omain-encoding genes (HRJDs) are a group of such genes (with typically two or three orthologs per species) characterized by their JmjC domain (Fig. 1A), yet their molecular functions remain unknown. Given their potential influence on the regenerative process, HRJDs may contribute to the high regeneration potential of highly regenerative animals. With this in mind, a question arises: what would happen if a low regenerative species, which has lost HRJDs, were to acquire them again? By ectopically expressing HRJDs in low regenerative animal models, we can investigate their impacts on regeneration as well as on aging processes, providing insight into the role of HRJDs.

**Figure 1.**
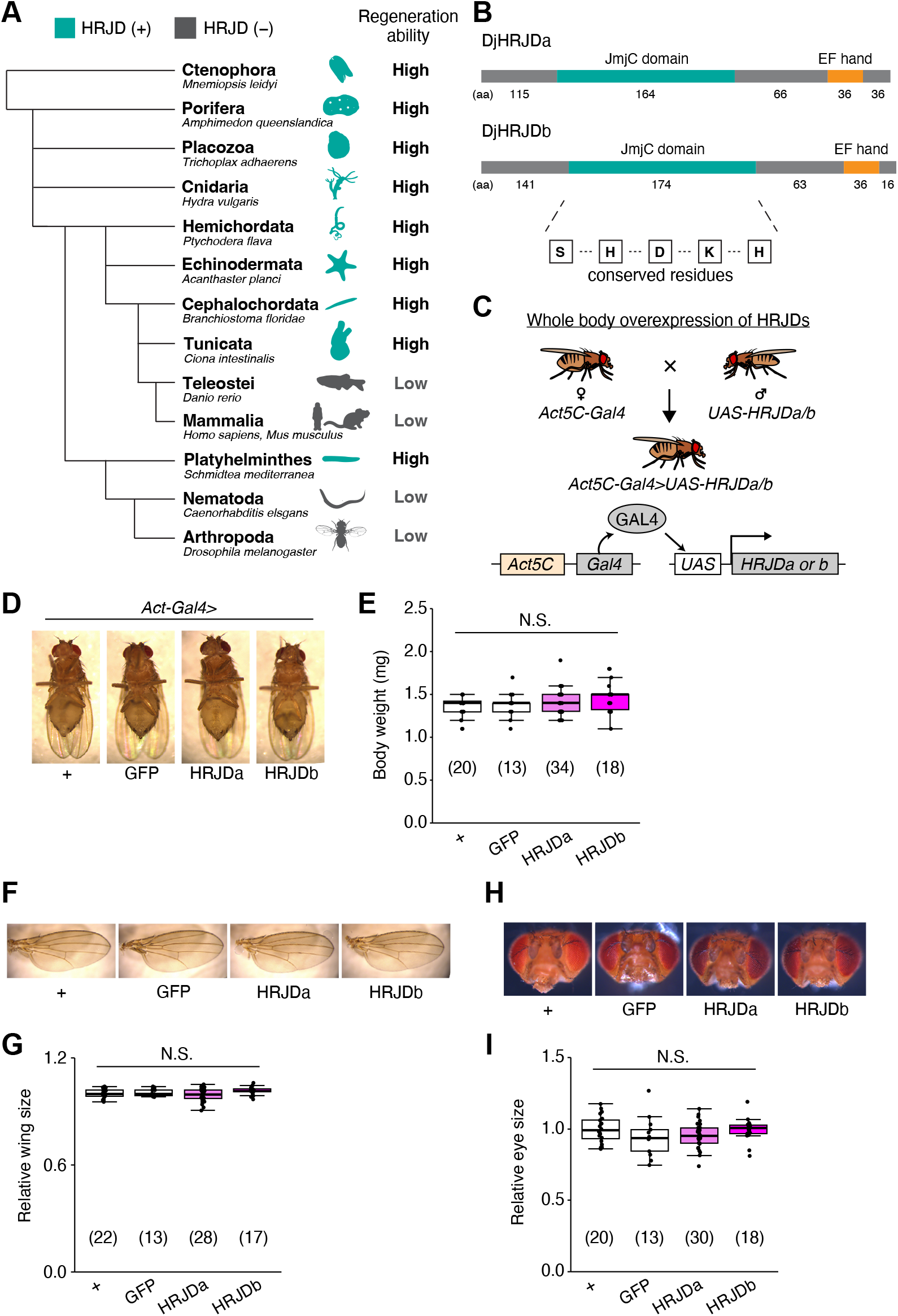
HRJD expression in whole body does not affect gross morphology. (A) Phylogenetic tree of HRJD conservation. Green indicates species that possess HRJD gene(s) and gray indicates species that have lost HRJD gene(s). High regenerative ability indicates that the species can regenerate their whole body or anterior/posterior body parts, and low regenerative ability indicates that the species can only regenerate their appendage (limbs, tails, fins) or much smaller scale of tissues/organs. We referred to Cao et al., 2019 for the definition of regenerative ability [13]. (B) Protein sequence of HRJDs used in this study, which derived from *Dugesia japonica* (DjHRJDa/b, hereafter simply described as HRJDa/b). Conserved residues characteristic of the JmjC domain are shown. (C) Schematics of genetic experiments for whole body induction of HRJDs. The Gal4-UAS system enables gene expression downstream of the UAS sequence, which is regulated by the transcription factor Gal4 [15]. In this case, the ubiquitously active *Act5C* promoter is used for Gal4 expression. (D) Representative images of mature adults for whole body expression of HRJDs. (E) Whole body induction of HRJDs did not change body weight of mature adults. (F) Representative images of adult wings. (G) Whole body induction of HRJDs did not change wing size. (H) Representative images for adult heads. (I) Whole body induction of HRJDs did not change eye size of adult flies. N.S., not significant: P>0.05. One-way ANOVAs with post hoc Tukey test. *n* indicates the number of flies examined. See also Fig. S1.

Here we express HRJDs in the fruit fly *Drosophila melanogaster* and evaluate their impact *in vivo*, especially by focusing on two epithelial tissues: developing wing discs and post-developmental adult midguts, both of which exhibit regeneration potential and can replenish damaged epithelial cells. In contrast to the predicted contribution of HRJDs in regeneration as observed in planarian, ectopic HRJD induction impedes regenerative responses and decreases organismal survival upon injury in *Drosophila*. Surprisingly, however, HRJD expression in the stem/progenitor population of the adult midguts extends organismal lifespan under the non-regenerative condition. Further investigations reveal that HRJDs enhance the proliferative activity of intestinal stem cells while keeping their differentiation fidelity in aged guts, ameliorating age-related decline in gut barrier functions. These findings provide evidence that genes specific to highly-regenerative animals can improve stem cell function as well as increase healthy lifespan upon heterologous expression in aging animals.

## RESULTS

### Planarian HRJDs expression do not affect gross morphology of fly adults

Planarians are one of the most highly regenerative animals; they are capable of regenerating most body parts upon amputation and can even reconstruct their whole body from fragments [1–4, 8, 14]. Previous work has identified two HRJD orthologs *HRJDa* and *HRJDb* from two planarian species, *Dugesia japonica* and *Schmidtea mediterranea*, where both HRJDs contain only the JmjC domain and the EF hand motif (Fig. 1B) [13]. In functional assays using RNAi-mediated knockdown in *D. japonica*, these two HRJDs affect viability of organisms after amputation [13], suggesting that planarian HRJDs are associated with regeneration processes. We thus utilized these functional HRJDs in *D. japonica* as representatives for the following studies and hereafter simply named them as HRJDa and HRJDb.

To examine potential benefits and/or disadvantages of acquiring HRJDs, we then ectopically expressed HRJDs in *Drosophila melanogaster*, which has lost HRJD genes during evolution (Fig. 1A), using the Gal4/UAS system (Fig. S1) [15]. We first introduced HRJDs in the whole body throughout development with the ubiquitous driver *Act5C-Gal4* (Fig. 1C, *Act5C-Gal4>UAS-HRJDa/b*). The whole body expression of HRJDs neither caused developmental lethality nor changed the body weight of mature adults compared with the *Act5C-Gal4>UAS-GFP* control (Fig. 1D and 1E). We further assessed the gross morphology of adult wings and eyes under the ubiquitous expression of HRJDs. The wing size was not altered by HRJD expression (Fig. 1F and 1G). Similarly, *Act5C-Gal4>UAS-HRJDa/b* flies did not change the size of adult compound eyes (Fig. 1H and 1I). These results indicate that HRJD expression does not disturb gross morphology of adult flies under homeostatic conditions, likely due to minimal impacts on developmental processes.

### HRJD expression hampers tissue regeneration in the developing wing disc

Given that HRJDs are conserved only among highly regenerative animals, their primary functions may be related to regeneration processes. Indeed, both planarian HRJDa and HRJDb function in whole-body regeneration while their relative contribution is likely context-dependent: HRJDa is indispensable for the regeneration of amputated heads while HRJDb promotes organismal survival after two consecutive amputations [13]. To test whether heterologously-induced HRJDs can enhance regenerative responses in *Drosophila*, we examined their impacts on regeneration after ablation of the developing wing imaginal disc. The *Drosophila* larval imaginal discs, including wing discs, which are composed of columnar epithelial cells, exhibit regenerative capacity and restore morphology even after massive cell death [16, 17] (Fig. 2A and 2B). We utilized the genetic ablation system in which transient overexpression of *eiger*, a *Drosophil*a TNF ligand, induces apoptosis in the wing pouch region [17, 18]. In this tissue ablation system, the temperature-sensitive form of the Gal4 repressor Gal80 (*tub-Gal80^ts^*) allows transient tissue ablation under the control of a wing pouch driver *rn-Gal4.* To circumvent the temporal expression associated with the Gal4/Gal80^ts^ system, we further introduced an additional binary expression system: the QF/QUAS system for HRJD induction (Fig. 2A) [19, 20]. When expressing HRJDs in the wing pouch with the *WP-QF* driver during the entire process of recovery, we found that *WP-QF>QUAS-HRJDs* flies exhibit severe defects compared to controls (Fig. 2C). Immunostaining of a mitotic marker, phospho-histone H3 (PH3), revealed that HRJD expression resulted in a slight but not statistically significant decrease in regenerative proliferation in the wing pouch, implying the possibility of insufficient damaged cell replenishment (Fig. 2D and 2E). Importantly, HRJD induction in the wing disc affected neither the development of the adult wing nor proliferation in wing discs under homeostatic condition (Fig. 2F-2I). These results suggest that HRJD expression does not facilitate regeneration in the developing wing disc epithelium.

**Figure 2.**
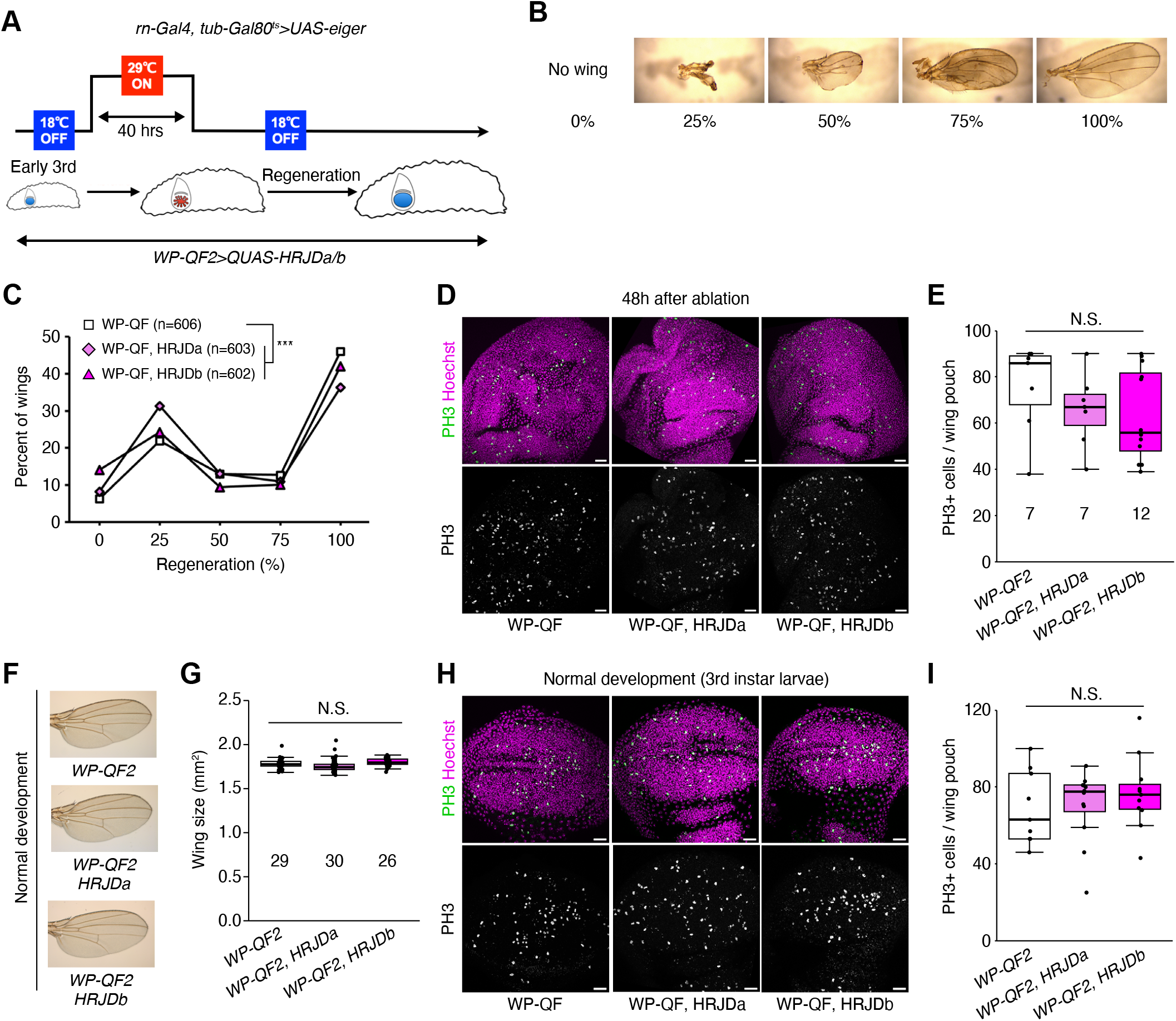
HRJD expression impedes tissue regeneration in the developing wind disc. (A) Schematics of genetic ablation of the wing disc. TNF ligand *eiger* was overexpressed in the wing pouch for 40 hrs during the larval stage. (B) Wing regeneration was assessed in the adult stage by checking the size of wing (0%: no wing, 100%: intact wing). (C) Line graph for wing regeneration. Expression of HRJDa/b increased the rate of low regeneration (0% and 25%) at the expense of the rate of full regeneration (100%). (D-E) Representative images for PH3 staining of wing discs during regeneration at 48 hrs after ablation (D). The number of PH3 positive cells is quantified in (E). (F-G) Representative images for adult wings of *WP-QF>QUAS-HRJDa/b* flies under homeostatic (non-regenerative) condition (F). The wing size is quantified in (G). (H-I) Representative images for PH3 staining of wing discs (3rd instar larvae) during normal development (H). The number of PH3 positive cells is quantified in (I). N.S., not significant: P>0.05, ***P ≤0.001. Chi-square test (C) and one-way ANOVAs with post hoc Tukey test (E, G, I). *n* indicates the number of wings (C, G) and wing discs (E, I) examined. Scale bars: 20 µm.

### HRJD expression compromises regeneration in the adult midgut

Most highly regenerative animals exhibit regeneration potential in mature adult stages [2–4, 8], and thus one possibility is that HRJDs stimulate regenerative responses only in adults, where tissue-resident stem cells play an important role in homeostasis and regeneration. We thus tested the impact of HRJD expression on the regenerative capacity of the *Drosophila* adult midgut. In the adult midgut, intestinal stem cells (ISCs) self-renew ISCs themselves and also generate differentiating progenitor cells called enteroblasts (EBs) and enteroendocrine progenitors (EEPs), which eventually differentiate into absorptive enterocytes (ECs) and secretory enteroendocrine cells (EEs), respectively (Fig. 3A) [21–24]. The adult midgut activates proliferation of ISCs in response to orally-treated harmful chemicals such as paraquat and dextran sulfate sodium (DSS), a regenerative response that is essential for organismal survival [21, 25–28]. We thus expressed HRJDs in ISCs and EBs using the *esg-Gal4* (*esg-Gal4>UAS-HRJDa/b*) driver and tested survival against paraquat/DSS damage. Similar to the wing regeneration assay, continuous expression of HRJDs via *esg-Gal4* significantly impaired organismal survival upon the chemical challenges, with a stronger effect for HRJDa (Fig. S2A and S2B). Survival under control feeding (5% sucrose) was also slightly decreased by HRJD expression, albeit not to a statistically significant degree (Fig. S2C). We further examined ISC proliferation after paraquat feeding by counting the number of cells marked with a mitotic marker, anti-PH3 staining, and found that induction of HRJDa significantly suppressed ISC division (Fig. S2D), which is consistent with the stronger decline in survival rate. Concordant with the wing ablation experiments, these results suggest that continuous HRJD expression causes detrimental effects to regenerative capacity.

**Figure 3.**
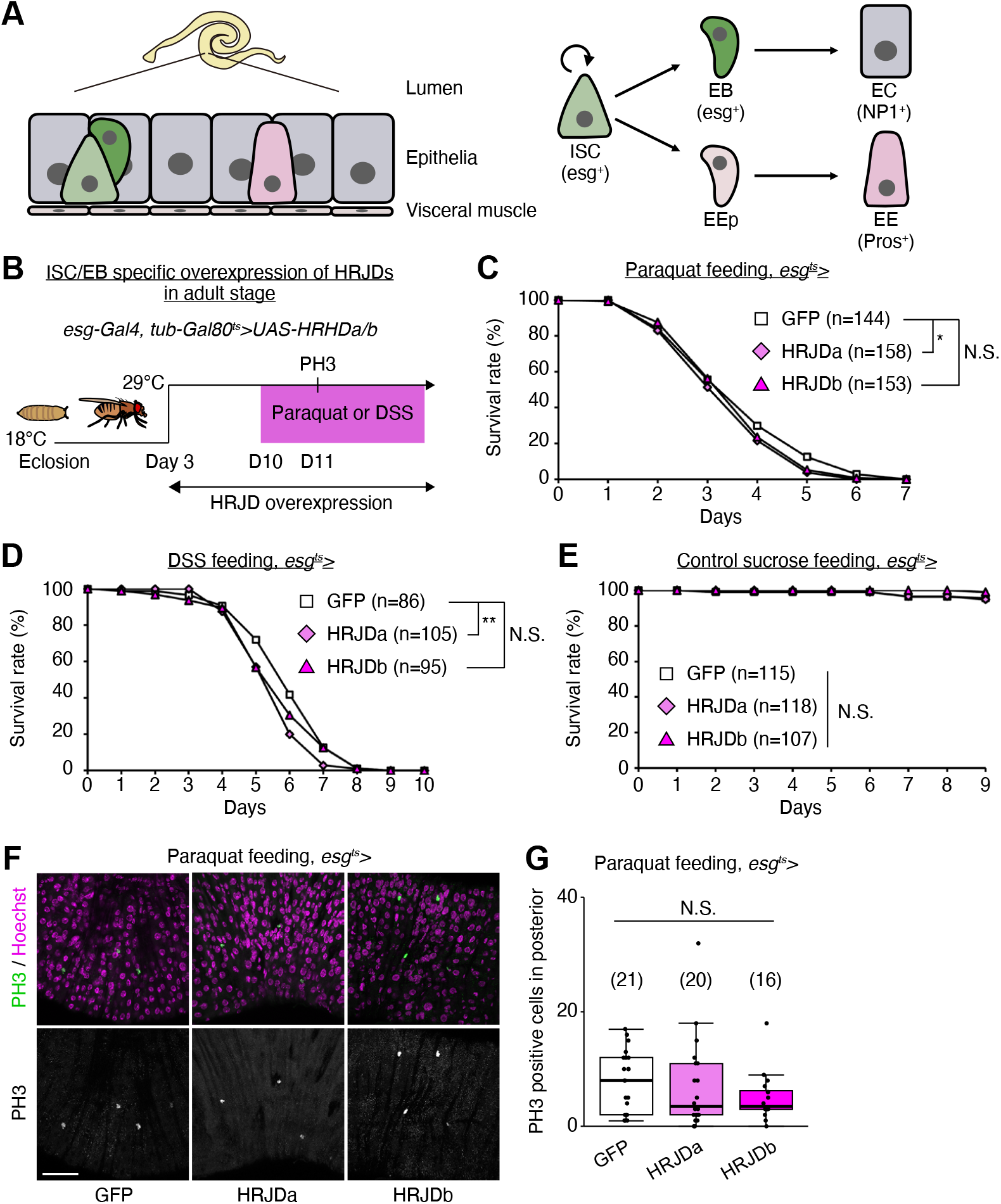
HRJDs expression compromises regeneration in the adult midgut. (A) Schematics of the *Drosophila* adult midgut. The adult midgut is a pseudostratified epithelium, in which intestinal stem cells (ISCs) generate both absorptive enterocytes (ECs) and secretory enteroendocrine cells (EEs) through progenitor cell enteroblasts (EBs) and EE progenitor cells (EEPs). (B) Experimental scheme for ISC/EB specific induction of HRJDs under the *esg-Gal4* driver in the adult stage. Adult flies were transferred to 29°C at Day 3 to induce HRJD expression. (C-E) Survival curve during paraquat (C), DSS (D), and sucrose (E) feeding. (F) Representative images for PH3 staining upon paraquat feeding. (G) Quantification of PH3 positive cells in posterior midguts. N.S., not significant: P>0.05, *P<0.05, **P≤0.01. Log-rank test. *n* indicates the number of flies (C-E) and midguts (G). Scale bars: 50 µm. See also Fig. S2.

Because the *esg-Gal4* driver is active not only in adult ISCs/EBs but also in the proliferative adult midgut progenitors in the larval midguts and larval imaginal discs [29, 30], continuous HRJD expression may cause internal organ defects associated with development. To temporally restrict HRJD expression to the adult stage, we next set up an experiment where HRJD expression starts only in adults after eclosion from pupa by combining *esg-Gal4* with *tub-Gal80^ts^* (*esg-Gal4^ts^>UAS-HRJDa/b*) (Fig. 3B); however, neither HRJDa nor HRJDb improved organismal survival upon paraquat/DSS feeding (Fig. 3C and 3D). Instead, HRJDa expression caused a slight but significant decrease in survival rate compared to controls (*esg-Gal4^ts^>UAS-GFP*). HRJDb expression similarly resulted in decreased survival at late phase (Day 4 and Day 5 in paraquat feeding, Day 5 and Day 6 in DSS feeding), though not to a statistically significant degree (Fig. 3C and 3D). Survival under control feeding (5% sucrose) was comparable between *esg-Gal4^ts^>UAS-GFP* and *esg-Gal4^ts^>UAS-HRJDa/b* (Fig. 3E), suggesting that the impaired survival of *esg-Gal4^ts^>UAS-HRJDs* is specific to regenerative contexts. We also found that, after paraquat feeding, mitotic cell numbers in HRJD expressing flies decreased slightly (Fig. 3F and 3G), suggesting the possibility of impaired regeneration. These results indicate that post-developmental expression of HRJDs does not enhance intestinal regeneration in adult flies.

### Post-developmental HRJD expression in ISC lineages extends organismal lifespan

Highly-regenerative animals are often resistant to organismal aging and have long lifespan; some species like the jellyfish *Turritopsis* even revert their early developmental stages under harsh environments and are considered to be potentially immortal [2, 6]. Although heterologous expression of HRJDs does not augment tissue regeneration in either developing or adult *Drosophila* tissues, we further investigated whether HRJDs influence organismal lifespan under homeostatic conditions in which no experimental injuries are applied to fly adults. We observed that post-developmental HRJD expression in adult ISCs/EBs (*esg-Gal4^ts^>UAS-HRJDa/b*) significantly extended organismal lifespan both in females (Fig. 4A) and males (Fig. 4B). In contrast, continuous HRJD expression in the whole body (*Act5C-Gal4>UAS-HRJDa/b*) shortened males’ lifespan (Fig. 4C and 4D). Moreover, HRJD expression in differentiated ECs (*NP1-Gal4>UAS-HRJDa/b*) also negatively impacted female lifespan, likely due to developmental abnormalities since EC-specific HRJD expression in the adult stage did not decrease survival (Fig. S3A-S3D). These results suggest that spatio-temporally regulated induction of HRJDs in adult ISCs/EBs can be beneficial for adult flies under homeostatic conditions.

Further investigation revealed the distinct impacts of HRJDa and HRJDb on organismal lifespan: while continuous expression of HRJDa via *esg-Gal4 (esg-Gal4>UAS-HRJDa)* shortened lifespans for both males and females, *esg-Gal4>UAS-HRJDb* prolonged female lifespans but shortened male lifespans (Fig. S3E and S3F). The ortholog-dependent phenotypic differences were also reported in planarian regeneration [13]. To address the mechanisms of lifespan extension associated with post-developmental HRJD expression in ISCs/EBs, we focused on *esg-Gal4^ts^>UAS-HRJDs* midguts in the following experiments.

**Figure 4.**
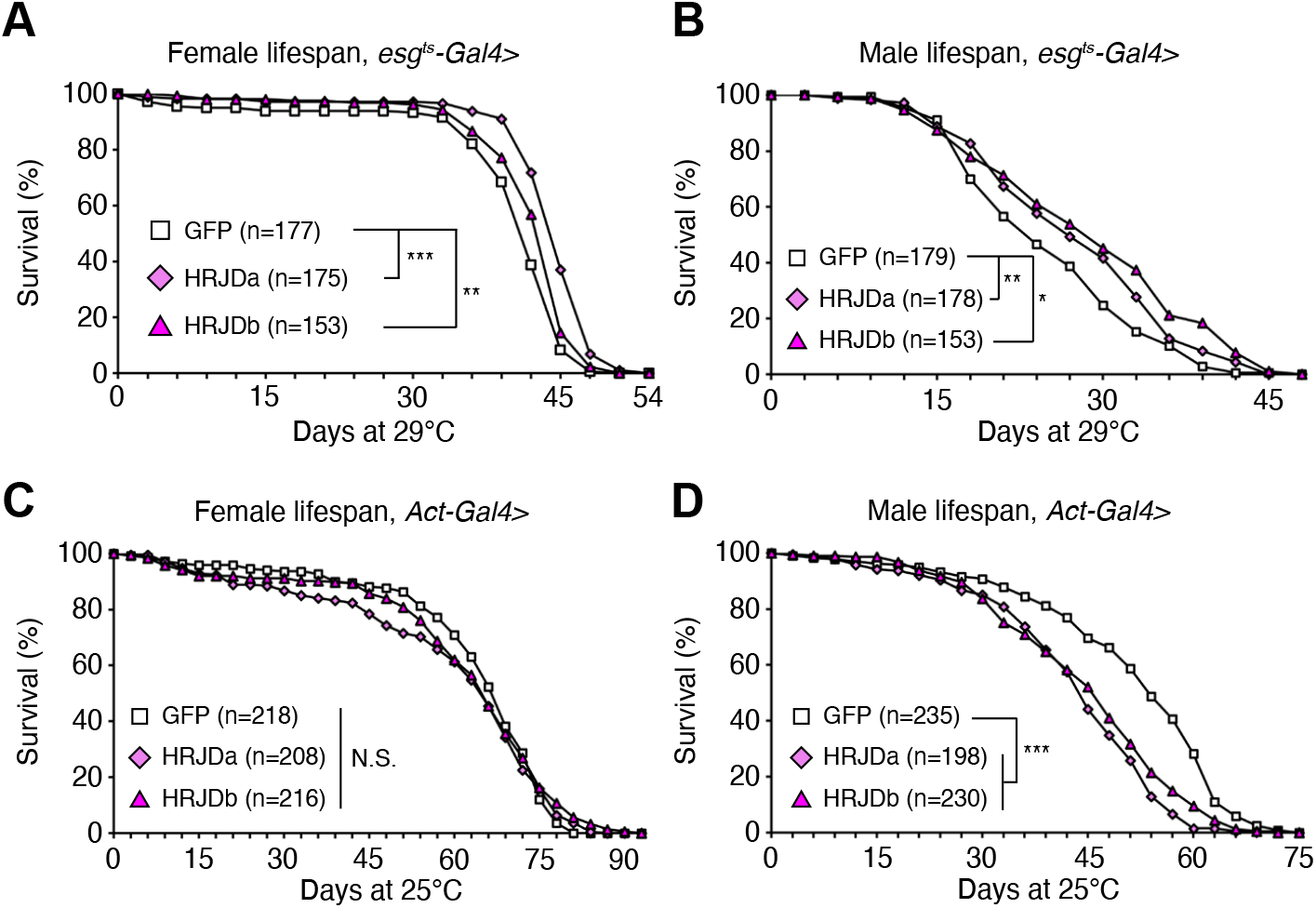
Post-developmental HRJD expression in ISC lineages extends organismal lifespan. (A-D) Survival curve for organismal lifespan by sex. HRJDs are induced by *esg^ts^-Gal4* (A and B) and *Act5C-Gal4* (C and D). For temporal regulation of HRJD induction using *tub-Gal80^ts^*(A and B), adult flies were transferred to 29°C at Day 3. N.S., not significant: P>0.05, *P<0.05, **P≤0.01, ***P≤0.001, Log-rank tests. *N* indicates the number of flies. See also Fig. S3.

### HRJD expression improves barrier function in the aged intestine

Age-related mortality accompanies intestinal barrier dysfunction and disruption of cell-cell junction in the gut epithelium across species [31–34]. Given that aging phenotypes are more severe in female flies than male flies [35], we first tested barrier function of the *esg-Gal4^ts^>UAS-HRJDa/b* female midgut using the Smurf assay, in which orally-administered blue dye spreads throughout the entire body when the intestinal barrier is disrupted (Smurf+, Fig. 5A) [31, 35]. Consistent with lifespan extension, post-developmental HRJD expression in ISCs/EBs significantly decreased the ratio of Smurf+ adults at 30-days old, a time point where age-related organismal death starts (Fig. 4A and 5A), suggesting that barrier function is maintained in aged flies that express HRJDs in ISCs/EBs. We further assessed the localization of septate junction markers (Dlg, Tsp2A, Mesh, Ssk), which diffuse in cytoplasm in the aged midgut [33, 36–38]. The diffusion of the Dlg protein in aged midguts was significantly suppressed in *esg-Gal4^ts^>UAS-HRJDa/b* midguts (Fig. 5B and 5C). Similarly, peripheral localization of Tsp2A, Mesh, and Ssk proteins was also maintained in *esg-Gal4^ts^>UAS-HRJDa/b* midguts compared with control (*esg-Gal4^ts^>UAS-GFP*) midguts (Fig. 5D-5I). Given that HRJDs in ISCs/EBs affect localization of junctional components in ECs, we next tested if HRJDs non-autonomously affect junctional integrity. To this end, we performed a mosaic experiment where HRJDs were clonally induced using the esgFLPout system [26]. Clonal induction of HRJDs improved junctional localization of Ssk both inside and outside the clones (Fig. S4A and S4B), supporting the conclusion that HRJDs’ effect is non-cell autonomous. We then examined JNK activation in ECs, which is associated with barrier dysfunction in the aged intestine [27, 36]. HRJD expression in ISCs/EBs significantly repressed transcription of *puckered* (*puc*), a downstream target of JNK signaling in the whole midgut (Fig. 5J), and indeed the intensity of *puc-lacZ* reporter decreased in ECs upon HRJD induction (Fig. 5K and 5L). HRJD expression in ISCs/EBs thus suppresses JNK signaling in ECs in the aged midgut, which is mediated by either a non-autonomous function of HRJDs or residual HRJD proteins in newly differentiated ECs. These results indicate that HRJD expression in adult ISCs/EBs contributes to the prolonged maintenance of the junctional integrity as well as gut barrier function in the aged intestine.

**Figure 5.**
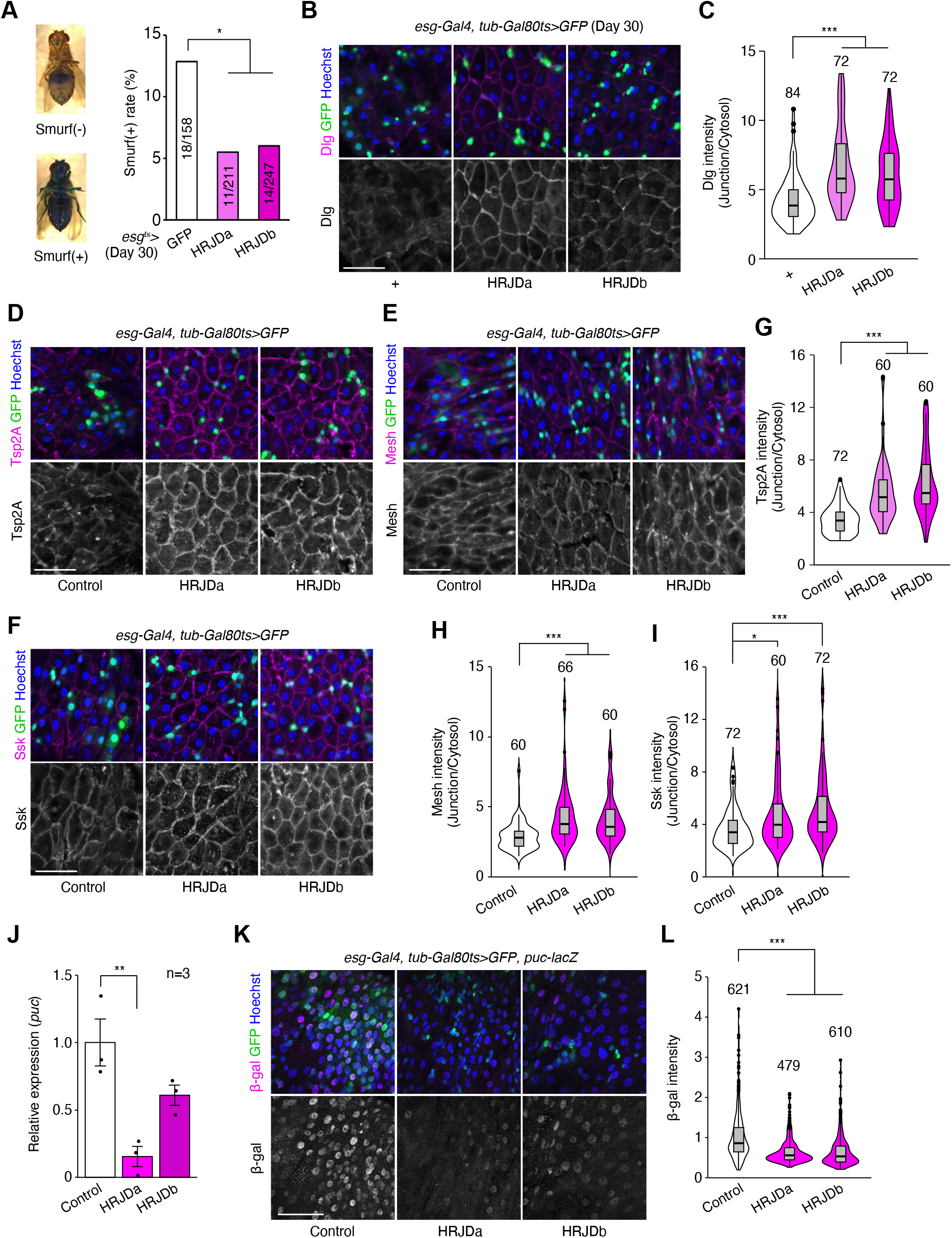
HRJD expression in adult ISCs/EBs suppress age-related gut barrier dysfunction. (A) *esg^ts^-*mediated HRJD induction decreased Smurf(+) ratio at Day 30. The representative images of the Smurf phenotype (leakage of blue dye throughout whole body) is shown on the left. (B-C) HRJD expression in adult ISCs/EBs maintained junctional localization of Dlg protein in ECs, which is quantified in (C). (D-I) HRJD expression in adult ISCs/EBs maintained junctional localization of Tsp2A (D), Mesh (E), and Ssk (F) in ECs, which is quantified in (G-I). (J) RT-qPCR of *puc* in the whole midgut. *esg^ts^-*mediated induction of HRJDa significantly repressed *puc*. (K-L) *puc-lacZ* intensity in polyploid cells was decreased by HRJD expression (K), which is quantified in (L). *P<0.05, **P<0.01, ***P≤0.001, One-way ANOVAs with post hoc Tukey test. *n* indicates the number of flies (A), cells (C, G, H, I, L), and biological replicates (J). Scale bars: 50 µm.

### HRJD expression attenuates mis-differentiation of stem cell lineage in the aged intestine

In addition to the intestinal barrier dysfunction, aberrant activation of ISC mitosis and accumulation of mis-differentiated ISC progenies are also common age-related phenotypes in the *Drosophila* adult midgut [27, 31, 33, 39]. In 30-days old control midguts, *esg-Gal4^ts^>UAS-GFP^+^*cells exhibited hallmarks of age-related mis-differentiation into ECs such as increased ploidy and enlarged nuclear size (Fig. 6A) [27]. By contrast, we found that HRJD expression in adult ISCs/EBs suppressed these mis-differentiation phenotypes (Fig. 6A and 6B), which are consistent with the amelioration of age-related barrier dysfunction in *esg-Gal4^ts^>UAS-HRJDa/b* midguts. Surprisingly, however, *esg-Gal4^ts^>UAS-HRJDa/b* did not suppress age-related increase in ISC proliferation, but rather enhanced it in 30-day old midguts (Fig. 6C and 6D). The enhancement of ISC proliferation was specific to the aged intestine since *esg-Gal4^ts^>UAS-HRJDs* did not affect PH3^+^ cell number in 10-day-old young midguts (Fig. S4E). These results raised the possibility that HRJD expression augments the mitotic activity of stem cells while maintaining their differentiation fidelity during aging.

**Figure 6.**
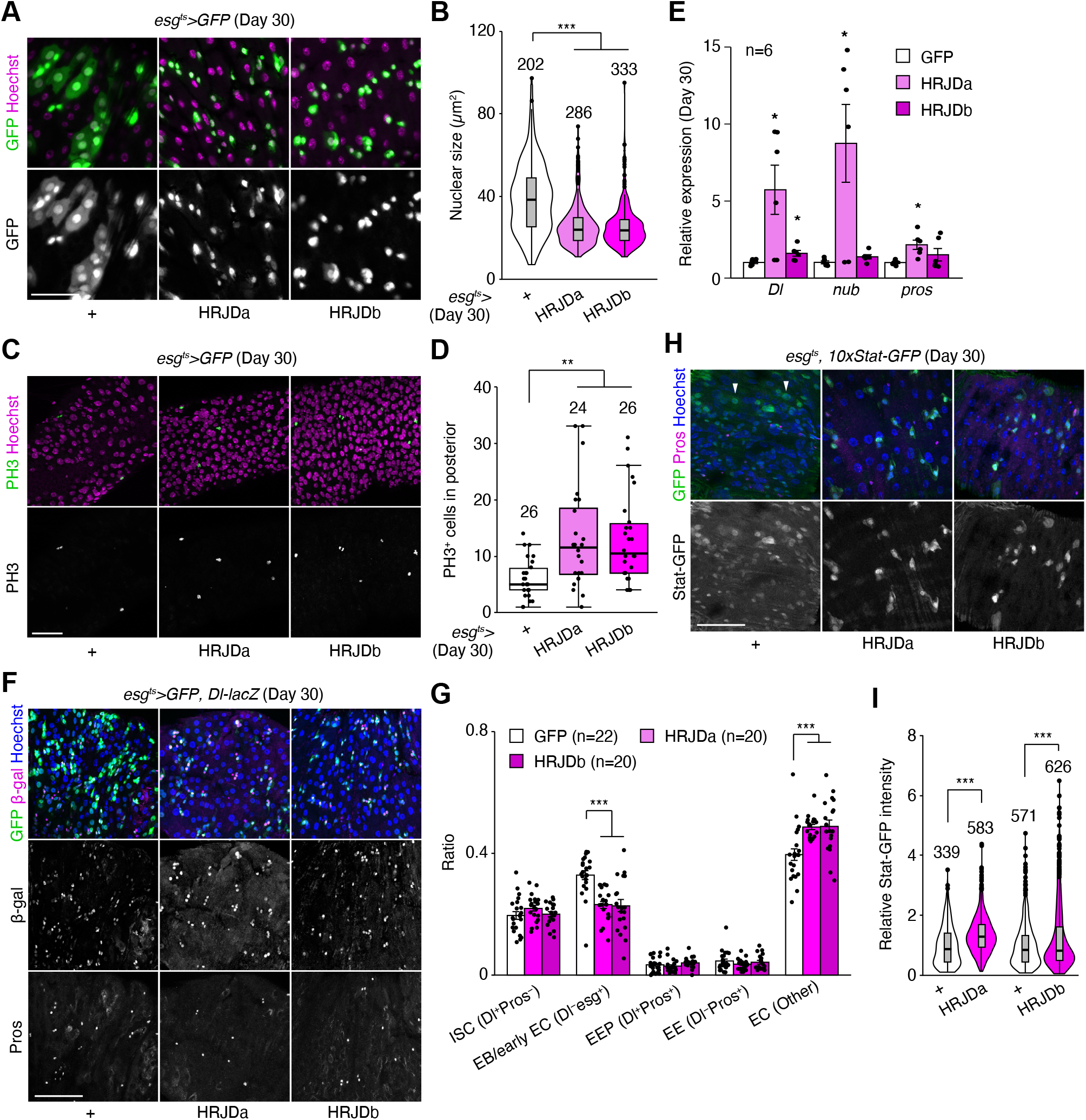
HRJD expression in adult ISCs/EBs improves stem cell functions in aged midguts. (A) Representative images for *esg-Gal4* positive cells in Day 30 midguts. HRJDs suppressed mis-differentiation of *esg^+^*cells (cellular enlargement with polyploid large nuclei). (B) Quantification of nuclear size of *esg-Gal4* positive cells. (C-D) Representative images (C) and quantification (D) of PH3 positive cells in Day 30 posterior midguts. (E) RT-qPCR of *Dl*, *nub*, and *pros* in Day 30 midguts. Expression levels were normalized to those of GFP control. (F-G) Cell type composition in aged midguts were assessed with *esg^ts^>GFP* (ISC, EB, and early EC), *Dl-lacZ* (ISC and EEP), and anti-Pros staining (EEP and EE). HRJD expression in adult ISCs/EBs reduced the ratio of EB/early EC and increased the ratio of EC (G). (H-I) *esg^ts^*-mediated HRJD induction enhanced STAT-GFP intensity in Pros^−^ diploid cells (ISCs/EBs), which is quantified in (I). Additionally, HRJDs suppressed STAT activation in polyploid ECs (arrowheads). N.S., not significant: P>0.05, *P≤0.05, **P≤0.01, ***P≤0.001, one-way ANOVAs with post hoc Tukey test. *n* indicates the number of cells (B, I), guts (D, G), and biological replicates (E). Scale bars: 50 µm.

To test this hypothesis, we first measured the expression of cell type markers (*Delta* for ISCs, *nub* for ECs, and *pros* for EEs) by RT-qPCR of whole midguts. Both *nub* and *pros* are negatively regulated by Esg [40, 41], and accumulation of *esg^+^* mis-differentiated cells accompanies the down-regulation of EC-related genes in aged midguts [27]. Consistent with enhanced ISC proliferation, *esg-Gal4^ts^>UAS-HRJDa* significantly up-regulated *Delta* (*Dl*) expression in 30-day old midguts (Fig. 6E). More importantly, *esg-Gal4^ts^>UAS-HRJDa* also up-regulated *nub* and *pros*, which supports our hypothesis that HRJDs improve proper differentiation of ISC progenies. In *esg-Gal4^ts^>UAS-HRJDb*, only *Dl*, but not *nub* and *pros*, was up-regulated when whole midguts were used as sample (Fig. 6E). Next, we focused on the posterior midgut where both HRJDa and HRJDb prevent mis-differentiation of EBs (Fig. 6A), and examined cell type composition using the combination of markers *esg^ts^>GFP* (ISC/EB), *Dl-lacZ* (ISC/EEP), and anti-Pros (EEP/EE). Consistent with the prevention of EB mis-differentiation, *esg^ts^*-mediated HRJD expression decreased EBs (esg^+^Dl^−^Pros^−^) and increased polyploid ECs (esg^−^Dl^−^Pros^−^) compared to the control midgut (Fig. 6F and 6G). These results indicate that HRJDs improve differentiation fidelity in ISC lineage.

To address the potential mechanism of HRJD-dependent maintenance of ISC functions, we focused on the JAK-STAT pathway since its activation promotes both ISC proliferation and differentiation into ECs [26, 42, 43]. In young mature midguts, STAT activity is largely restricted to *esg^+^* ISCs/EBs [26, 42, 44]. In aged midguts, however, a subset of polyploid cells exhibited weak signal of the STAT reporter *10×STAT-GFP*, likely due to the accumulation of mis-differentiated EBs (Fig. 6H). Notably, HRJD induction canceled such ectopic STAT activation in polyploid cells and rather enhanced STAT activity in diploid ISCs/EBs (Fig. 6H and 6I), which is consistent with our observations that HRJDs boost ISC functions during aging. These results suggest that HRJD expression accelerates the generation of differentiated ECs via JAK-STAT activation. In the healthy homeostatic intestine, ISC division and subsequent differentiation into ECs is coupled with the loss of old ECs [45, 46]. Interestingly, we found that *esg*-lineage clones that express HRJDs rapidly replaced pre-existing cells by generating polyploid ECs (Fig. S4A, S4C, and S4D). In addition, more cells exhibited cleaved Dcp1, an apoptotic marker, in 30-day old *esg^ts^ >HRJDs* midguts (Fig. S4F and S4G), implying that HRJD induction promotes turnover of midgut epithelial cells. Collectively, our data show that post-developmental expression of HRJDs improves stem cell functions in the aged intestine.

## DISCUSSION

In this study, we demonstrate that post-developmental expression of planarian HRJD genes in the *Drosophila* adult ISCs/EBs can suppress age-related intestinal dysfunctions and extend organismal lifespan, while their continuous expression throughout the entire developmental process hampers regenerative responses and principally shortens lifespan. Notably, HRJDs in adult ISCs/EBs boost the age-related increase of ISC proliferation but do not cause age-related mis-differentiation of ISC progenies (Fig. 7). These HRJD-mediated outcomes are distinct from those mediated by typical anti-aging manipulations such as antibiotic treatment and metabolic intervention, which ameliorate both ISC over-proliferation and mis-differentiation [24, 31, 47]. Therefore, heterologously induced HRJDs create the neomorphic state in the aged intestine (Fig. 7). Given the age-related increase of chronic cellular stresses [24, 27, 36, 47, 48], we speculate that the HRJD-dependent ISC activation and their proper differentiation improve organismal fitness by enabling active turnover of damaged intestinal cells.

**Figure 7.**
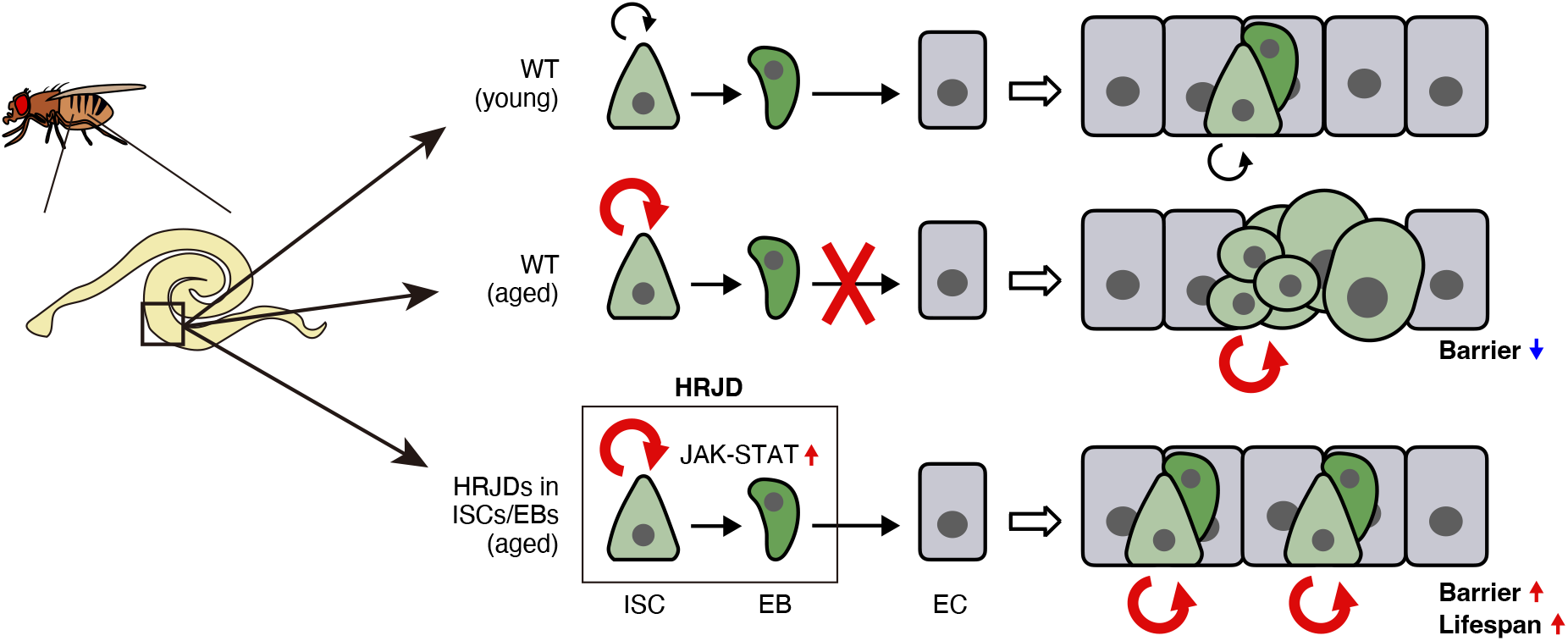
Graphical summary. Schematic model for the impact of HRJDs on intestinal homeostasis. In aged wildtype midguts, ISCs over-proliferate, and their daughters fail to differentiate into mature ECs, resulting in an accumulation of mis-differentiated cells (*esg*^+^ large polyploid). HRJD expression in adult ISCs/EBs further enhances ISC proliferation but suppresses mis-differentiation likely through upregulation of JAK-STAT signaling, which results in successful maintenance of the gut barrier and an extension of organismal lifespan.

Our investigation implies that the activation of the JAK-STAT pathway underlies the HRJD-mediated enhancement of ISC functions. Surprisingly, however, the expression of most mitogens including upstream ligands of JAK-STAT signaling (*upd1, upd2, upd3*) were not up-regulated by HRJD induction (Fig. S4H), raising the possibility that HRJDs activate STAT by modulating intracellular signaling factors. Such ligand-independent STAT activation can be achieved via non-receptor type tyrosine kinases like Src and Abl [49]. Given that the typical JmjC family proteins function as histone demethylase or protein hydroxylase [50], HRJDs may epigenetically target these tyrosine kinases. Notably, *Drosophila* Jarid2 (Jumonji, AT rich interactive domain 2) activates EGFR signaling upon overexpression in ISCs/EBs without changing ligand expression [51]. Our findings, together with these reports, suggest a cell-autonomous role of JmjC family proteins in regulating proliferative signaling in stem cell lineages. Intriguingly, Jarid2 expression in ISCs/EBs leads to ISC over-proliferation as well as barrier dysfunction, resulting in reduced lifespan [51], which is the opposite adult phenotype of HRJD expression in ISC lineages. Moreover, in contrast to the nuclear localization of histone demethylase KDM8 [52, 53], the closest paralog of HRJDs, ectopically-induced HRJDs localize in the cytoplasm but not in the nucleus both *in vitro* (S2 cells) and *in vivo* (wing discs and adult midguts) (Fig. S1B-S1D). This is a common localization pattern of JmjC-domain only proteins [53, 54] and implies that an unknown mechanism may operate when HRJDs work as a potential histone demethylase. In the future, it will thus be critical to examine the detailed molecular function of HRJDs to understand their impact on stem cell lineages.

Although HRJDs have been identified as genes conserved between highly regenerative animals [13], it is unclear whether HRJDs alone are sufficient to enhance regenerative responses. Our investigations revealed that ectopic expression of HRJDs failed to improve regeneration of developing wing discs and adult midguts in *Drosophila* (Fig. 2 and 3), likely through the attenuation of regenerative growth. However, it should be noted that we induced HRJDs before tissue injury and maintained their expression during regeneration period. Given that *HRJDb* is up-regulated in the late phase of planarian regeneration (3 days after amputation) [13], strict regulation of HRJD induction might be important for their proper function as true regeneration regulators. On the other hand, the expression level of *HRJDa* remains constant during planarian regeneration [13], which is recapitulated in our experiments. Another possibility is that a high regeneration ability can be achieved by cooperation between HRJDs and other unidentified genes that have been lost during evolution, which should be addressed in future studies.

During the normal aging process in the *Drosophila* adult midgut, ISC over-proliferation is closely linked with other aging phenotypes, and suppression of ISC division can prevent the accumulation of mis-differentiated ISC progenies and extend organismal lifespan [24, 36, 47, 48]. In contrast, HRJDs enhance age-related ISC activation but can alleviate mis-differentiation and extend lifespan, suggesting that it is not ISC activation itself but rather mis-differentiation of ISC progenies that principally drives age-related mortality. Consistently, manipulation of mitotic spindle orientation, which affects the cell fate of daughter cells [55–57], can extend organismal lifespan without changing mitotic activity of ISCs [57]. Of note, in contrast to *Drosophila* ISCs which over-proliferate in aged midguts, many adult stem cells decrease their activity and abundance during aging both in *Drosophila* and mammals [24, 58, 59]. Future investigations for mechanisms of HRJD-mediated stem cell rejuvenation will provide clues to develop new anti-aging strategies.

## CONCLUSIONS

In this study, we established a *Drosophila* model in which HRJDs are heterologously expressed in specific tissue or cell type using binary expression systems, and demonstrated that HRJDs can improve proliferative/differentiation capacity of ISCs in the aged midgut. Although continuous HRJD expression in the wing disc and in the adult midgut impairs tissue regeneration upon injury, restricted HRJD expression in post-developmental adult ISCs/EBs enhances mitotic activity of ISCs as well as maintains their differentiation fidelity in the aged flies, leading to the prevention of age-related intestinal barrier dysfunction and the extension of organismal lifespan. Our HRJD-expressing model will serve as a valuable resource to understand unprecedented mechanisms of stem cell rejuvenation in the future.

## METHODS

### *Drosophila* stocks

All stocks were maintained on a standard diet containing 4% cornmeal, 6% baker’s yeast (Saf Yeast), 6% glucose (Wako, 049-31177), and 0.8% agar (Kishida chemical, 260-01705) with 0.3% propionic acid (Tokyo Chemical Industry, P0500) and 0.05% nipagin (Wako, 132-02635). Canton S was utilized as the wildtype strain. Transgenic fly lines were obtained from the Bloomington Drosophila Stock Center (BDSC) and the Kyoto Stock Center: *Act5C-Gal4* (BDSC 3954), *rn-Gal4, tub-Gal80ts, UAS-egr* (BDSC 51280), *WP-QF2* [60], *tub-Gal80ts* (BDSC 7017, 7019), *esg-Gal4* (Kyoto 109126), *NP1-Gal4* (Kyoto 112001), *UAS-GFP* (BDSC 1522), *M{3xP3-RFP.attP}ZH-86Fb* (Kyoto 130437), *UAS-HRJDa* (this study), *UAS-HRJDb* (this study), *QUAS-HRJDa* (this study), *QUAS-HRJDb* (this study), *UAS-FLAG-HRJDa* (this study), *UAS-FLAG-HRJDb* (this study), *puc-lacZ* (BDSC 98329), *10×STAT-GFP* (BDSC 26198), *Dl-lacZ* (BDSC 11651). *esg-Gal4, UAS-GFP, tub-Gal80ts; UAS-FLP, Act5C-FRT.CD2-Gal4* (esgFLPout) is a gift from Irene Miguel-Aliaga. See **Table S1** for genotypes in each Figure. We used female flies unless otherwise noted in the Figures.

### *Drosophila* genetics

Experimental crosses that did not involve Gal80^ts^-mediated inhibition of Gal4 were performed at 25°C. When using Gal80^ts^, experimental crosses were maintained at 18°C. For genetic wing ablation experiments (Fig. 2), F1 larvae were raised at 18°C until day 7, incubated at 29°C for the next 40 hr, and then maintained at 18°C until the adult hatched or dissection. For lifespan assays, midgut staining, and Smurf assays (Fig. 3-6); F1 adults were transferred to 29°C three days after eclosion until the experiments. Midgut staining and Smurf assays were performed after 7 or 27 days of 29°C incubation (young: Day 10 adults, old: Day 30 adults). For esgFLPout experiments, flies were maintained at 18°C until 50 days, and 50-day old adults were transferred to 29°C and analyzed after 10 days (final 60-day old).

### Generation of HRJD expressing lines

To express HRJDs using the Gal4/UAS or the QF/QUAS system, we constructed a vector containing *HRJDa* or *HRJDb* under the control of either the UAS sequence or QUAS sequence. Namely we amplified HRJDa and HRJDb sequence from codon-optimized synthesized DNAs (pUCIDT-HRJDa and pUCIDT-HRJDb, IDT, Supplementary texts S1) for *DjHRJDa* (Genbank # LC408963) and *DjHRJDb* (Genbank # LC408964), respectively. UAS-HRJDa and UAS-HRJDb were constructed by ligating HRJDa and HRJDb into pUAST-attB (DGRC 1419), respectively (digested by EcoRI/NotI). Similarly, QUAS-HRJDa and QUAS-HRJDb were constructed by ligating HRJDa and HRJDb into pQUAS-WALIUM20 (DGRC 1474), respectively (digested by NheI/EcoRI). To construct UAS-3xFLAG-HRJDs, we PCR-amplified the coding sequences of HRJDs from pUCIDT-HRJDs by gene-specific primers with 3xFLAG tag at 5’-end of the forward primers, and the amplicons were then ligated into a pUAST-attB vector digested by EcoRI/NotI using In-Fusion HD Cloning Kit (Takara, 639649). The landing site for each construct was {3xP3-RFP.attP}ZH-86Fb. Injection and selection were performed by WellGenetics (Taiwan, R.O.C.). Please see also Table S2 and Supplementary texts S1 for primer sequences and HRJDs sequences.

### S2 cell culture

*Drosophila* S2 cells were grown at 25°C in Schneider’s *Drosophila* medium (GIBCO, 21720001) supplemented with 10% (v/v) heat-inactivated fetal bovine serum (FBS), 100 U/mL penicillin, and 100 μg/mL streptomycin (FUJIFILM Wako, 168–23191).

### S2 cell immunostaining

To examine the expression pattern of HRJDs through immunostaining, 1 × 10^6^ cells were seeded in 6-well plates containing Schneider’s *Drosophila* medium supplemented with 10% (v/v) heat-inactivated fetal bovine serum, 100 U/mL penicillin, and 100 μg/mL streptomycin and were transfected with 800 ng of pAc5-3xFLAG-HRJDs using Effectene Transfection Reagent (QIAGEN, 301427) following the manufacturer’s protocol. To prepare pAc5-3xFLAG-HRJDs, we amplified the coding sequences of HRJDs from pUCIDT-HRJDs (Supplementary texts S1) by gene-specific primers with 3xFLAG tag at 5’-end of the forward primers, and the amplicons were then ligated into a pAc5-STABLE2-neo (Addgene, 32426) [61] digested with EcoRI/XhoI by In-Fusion HD Cloning Kit (Takara, 639649). Twenty-four hours after transfection, the cells were washed with PBS and the medium was replaced with fresh Schneider’s *Drosophila* medium supplemented with 10% (v/v) heat-inactivated FBS, 100 U/mL penicillin, and 100 μg/mL streptomycin. Forty-eight hours after transfection, the cells were fixed with 4% paraformaldehyde (PFA) in PBS, washed with PBS containing 0.1% Triton X-100 (PBST), blocked in PBST with 5% normal donkey serum (PBSTn), and incubated with primary antibodies in PBSTn overnight at 4 °C. The samples were then washed with PBST, incubated for 1 h at room temperature with secondary antibodies and Hoechst 33342 suspended in PBSTn and washed again with PBST. Images were captured using an LSM880 (Zeiss). The primary antibody used was a mouse anti-FLAG M2 monoclonal antibody (1 : 2000, Sigma, F1804). The secondary antibodies used were Goat anti-Mouse IgG2b Cross-Adsorbed Secondary Antibody, Alexa Fluor™ 555 (1:2000, Thermo Fisher Scientific, A-21147). Hoechst 33342 (0.4 µM; Invitrogen, H3570) was used for nuclear staining. Images were analyzed and edited using Fiji/ImageJ software (NIH).

### S2 cell western blotting

To examine the expression of HRJDs, 2.5 × 10^5^ cells were seeded in 24-well plates containing Schneider’s *Drosophila* medium supplemented with 10% (v/v) heat-inactivated FBS, 100 U/mL penicillin, and 100 μg/mL streptomycin and were transfected with 200 ng of pAc5-3xFLAG-HRJDs using Effectene Transfection Reagent (QIAGEN, 301427) following the manufacturer’s protocol. Twenty-four hours after transfection, the cells were washed with PBS and the medium was replaced with fresh Schneider’s *Drosophila* medium supplemented with 10% (v/v) heat-inactivated FBS, 100 U/mL penicillin, and 100 μg/mL streptomycin. Forty-eight hours after transfection, the cells were washed with PBS and lysed with 50 µL RIPA buffer supplemented with cOmplete ULTRA EDTA-free protease inhibitor cocktail (Roche, 05892953001). The lysate was sonicated and centrifuged at 20,000 g for 5 minutes. Protein concentration was determined by BCA assay. The supernatant was mixed with SDS, boiled at 95 °C for 5 minutes, and subjected to western blotting.

Proteins were separated using SDS-PAGE and transferred onto Immobilon-P PVDF membranes (Millipore, IPVH00010) for immunoblotting. Membranes were blocked with 4% skimmed milk diluted in 1×Tris buffered saline containing 0.1% Tween-20. Immunoblotting was performed using the below-mentioned antibodies, which were diluted with 4% skim milk. The signals were visualized using Immobilon Western Chemiluminescent HRP Substrate (Millipore, WBKLS0500) and FUSION SOLO. 7S. EDGE (Vilber-Lourmat). Contrast and brightness adjustments were applied using the Fiji/ImageJ software (NIH).

The primary antibody used was mouse anti-FLAG M2 monoclonal antibody (1 : 5000, Sigma, F1804). Mouse anti-alpha tubulin (DM1A) monoclonal antibody (1 : 5000, Sigma, T9026) was used as loading control. The secondary antibody used was HRP-conjugated goat/rabbit/donkey anti-mouse IgG (1 : 10000, Promega, W402B).

### Drug treatment

5 mM paraquat (Sigma, 856177) and 5% (w/v) DSS (MP Biomedicals, 160110) were dissolved in 5% (w/v) sucrose solution. Filter paper (Whatman 3MM) was soaked with 400 µl of these reagents and placed into empty vials. For histological analyses, flies were fed with the reagent solution for 1 day. For the survival assay, flies were transferred to new vials and dead flies (determined by immobility and showing no response to tapping) were counted every day. 5% sucrose was used for control feeding.

### Immunofluorescence

Samples were dissected in 1×PBS and fixed in 4% PFA for 20 minutes (wing discs) and 30-45 minutes (adult midgut) at room temperature (RT), respectively. The following primary antibodies were used with indicated dilution into 1×PBS containing 0.5% BSA and 0.1% Triton X-100: rabbit anti-PH3 (Millipore 06-570, 1:1000), mouse anti-FLAG (Sigma F1804, 1:1000), mouse anti-Dlg (DSHB 4F3, 1:100), rabbit anti-GFP (MBL 598, 1:500), rat anti-GFP (Nacalai tesque 04404-26, 1:400), rabbit anti-Tsp2A (Izumi et al., 2016, 1:1000) [62], rabbit anti-Mesh (Izumi et al., 2012, 1:1000) [63], rabbit anti-Ssk (Izumi et al., 2016, 1:1000) [62], chicken anti-β-galactosidase (Abcam ab9361, 1:500), anti-cDcp1 (Cell Signaling Technology 9578, 1:200). After overnight incubation with primary antibodies at 4°C, samples were incubated with fluorescent secondary antibodies (Jackson ImmunoResearch and Invitrogen, 1:500) for 1 hour at RT. Hoechst 33342 (Invitrogen, final concentration: 10 µg/ml) was used to visualize DNA. Wing discs were mounted as described previously [64, 65]. Samples were mounted in Slowfade Diamond (ThermoFisher, S36963) and imaged with confocal microscopy Zeiss LSM880 or Zeiss LSM980.

### Smurf assay

We referred to Rera et al. (2012) for the Smurf assay conditions [35]. To prepare the feeding medium, 100 µl of 50% (w/v) brilliant blue FCF (Wako, 027-12842, final 2.5%) and 100 µl of 5% (w/v) sucrose were added to a vial containing 2 ml of cornmeal-agar food. After mixing with a spatula, an Whatman 1 filter (1001-020) was put on the feeding medium. Flies were fed with this medium at 25°C for one day, after which they were transferred to a new vial containing cornmeal-agar food without blue dye to clean the epidermis. Two hours after the transfer, Smurf phenotype was checked. We classified the Smurf phenotype as blue dye leakage outside abdomen (thorax, head, legs).

### RT-qPCR

Total RNA was purified from 10-15 midguts using the ReliaPrep RNA Tissue Miniprep System (Promega). cDNA was made from 100 or 200 ng of RNA using PrimeScript RT Reagent Kit (TaKaRa). Quantitative PCR was performed using TB Green Premix Ex Taq II (TaKaRa) and the QuantStudio 6 Flex Real-Time PCR System (ThermoFisher). RpL32 was used as an internal control. Primer sequences are listed in Table S2.

### Phylogenetic tree

We created the phylogenetic tree of representative organisms using the NCBI taxonomy browser (https://www.ncbi.nlm.nih.gov/Taxonomy/CommonTree/wwwcmt.cgi). The species examined are *Amphimedon queenslandica* (Porifera), *Mnemiopsis leidyi* (Ctenophora), *Trichoplax adhaerens* (Placozoa), *Hydra vulgaris* (Cnidaria), *Ptychodera flava* (Hemichordata), *Acanthaster planci* (Echinodermata), *Branchiostoma floridae* (Cephalochordata), *Ciona intestinalis* (Tunicata), *Danio rerio* (Teleostei), *Homo sapiens*, *Mus musculus* (Mammalia), *Schmidtea mediterranea* (Platyhelminthes), *Caenorhabditis elegans* (Nematoda), and *Drosophila melanogaster* (Arthropoda). These species, except for *D. rerio, H. sapiens, M. musculus, C. elegans,* and *D. melanogaster,* are known to have HRJDs and exhibit high regeneration ability [13]. The pictures were downloaded from PhyloPic, and the color of some were changed from black to blue. Credits: Bennet McComish (*B. floridae*), Malio Kodis (*M. leidyi*), and Markus A. Grohme (*S. mediterranea*, https://www.phylopic.org/images/93f9611c-0cbd-4a14-92da-004a4521e21f/schmidtea-mediterranea).

### Quantification of PH3 positive cells

For wing discs, we counted PH3 positive cells in the pouch region, which develops into the adult wing. After binarization, the number of PH3 positive cells was counted using the Analyze Particles function in Fiji/ImageJ, and subsequently confirmed or corrected by visual review of the images. For adult midguts, the number of PH3 positive cells in the posterior midguts was manually counted.

### Quantification of junctional proteins

To quantify junctional and cytoplasmic intensity, a line that was orthogonal to one side of a cell was drawn, and then the plot profile was calculated for the line. We defined the junctional intensity as the highest value in that prolife, and defined cytoplasmic intensity as the value 20 pixels (∼2.6 µm) away from the highest value. Three lines were drawn for one cell and the average of the three was used to represent the junctional/cytoplasmic ratio for the cell. Six polyploid cells were randomly selected from each image for quantification.

### Quantification of reporter intensity

The signal intensity of *puc-lacZ* and *10×STAT-GFP* reporters was quantified using Fiji/ImageJ. The nuclei of large polyploid cells (for *puc-lacZ*) and Pros^−^ diploid cells (for *10×STAT-GFP*) were selected as ROIs using the polygon selection tool. The reporter intensity in each ROI was quantified using the Measure command.

### Quantification of nuclear size

Nuclei of *esg-Gal4>UAS-GFP* positive cells were traced by the polygon selection tool and added as ROI in Fiji/ImageJ. The size of each ROI was quantified using the Measure command.

### Quantification of cell number

Cell number was quantified using Fiji/ImageJ. To count the total cell number, Hoechst signal was processed as following: (1) despeckle, (2) binarization, (3) fill hole, (4) watershed, (5) analyze particle. The following cell number was manually counted using the cell counter function: *esg>GFP^+^*cells, *Dl-lacZ^+^* cells, Pros^+^ cells, and cDcp1^+^ cells.

### Statistics

Statistical analyses were performed using Excel and RStudio. Two tailed *t* tests were used for comparisons between two groups. One-way ANOVAs with post hoc Tukey tests were performed when comparing three or more groups. Log-rank tests were used for comparison of survival curve. Significance is indicated in the Fig.s as follows: *P≤0.05, **P≤0.01, ***P≤0.001, Not Significant (N.S.): P>0.05. Bar graphs show mean ± standard error. Boxplots show median (thick line in the box), first and third quartiles (bottom and top of the box), minimum value (lower whisker), and maximum value (upper whisker). Dots in bar graphs and boxplots indicate individual values. Violin plots portray the distribution of individual values.

## Abbreviations

i-cell: interstitial cell
HRJD: highly regenerative species-specific JmjC domain-encoding gene
PH3: phospho-histone H3
ISC: intestinal stem cell
EB: enteroblast
EEP: enteroendocrine progenitor
EC: enterocyte
EE: enteroendocrine cell
DSS: dextran sulfate sodium
puc: puckered
Dl: Delta
Jarid2: Jumonji, AT rich interactive domain 2

## DECLARATIONS

### Ethics approval and consent to participate

Not applicable.

### Consent for publication

Not applicable.

### Availability of data and materials

The datasets and *Drosophila* stocks used and/or analyzed during the current study are available from the corresponding author on reasonable request.

### Competing interests

The authors declare that they have no competing interests.

### Funding

This work was supported by JSPS/MEXT KAKENHI (grant numbers JP22J01430 to H.N., JP21H04774, JP23H04766, JP24H00567 to M.M., and JP17H06332, JP22H02762, JP23K18134, JP23H04696 to Y.N.), AMED-Aging (JP21gm5010001 to M.M.), AMED-PRIME (JP22gm6110025 to Y.N.), and Sadako O. Hirai Ban Award for Young Researchers (H.N.)

### Author contributions

Conceptualization: H.N., Y.N.

Investigation: H.N., Y.A., T.N., E.T., J.U., Y.N.

Methodology: H.N., Y.N.

Validation: H.N., Y.A., T.N., E.T., J.U., T.M., M.M., Y.N.

Data curation: H.N., T.M., Y.N.

Writing – original draft: H.N., J.U., T.M., Y.N.

Writing – review & editing: H.N., Y.A., T.N., J.U., T.M., M.M., Y.N.

Supervision: M.M., Y.N.

Funding acquisition: H.N., T.M., M.M., Y.N.

## Acknowledgements

We thank I. Miguel-Aliaga, N. Shinoda, M. Furuse, Y. Izumi, BDSC, Kyoto Stock Center, Transgenic RNAi Project (TRiP), and DSHB for fly stocks and reagents.

## Supplemental figure legends

**Figure S1. Expression and localization of HRJD proteins**

(A) Western blots of S2 cell samples upon transfection of 3×FLAG-HRJDa/b. Anti-FLAG was used to detect 3×FLAG-HRJDa/b proteins and anti-tubulin was used as the loading control. Expected molecular weight: 52.0 kDa for 3×FLAG-HRJDa, 52.9 kDa for 3×FLAG-HRJDb.

(B) Representative images of anti-FLAG staining using S2 cells transfected with 3×FLAG-HRJDa/b. Both HRJDa and HRJDb localized cytoplasm but not in nuclei. Scale bar: 30 µm in wide-view images (left), 5 µm in magnified view images (yellow frames). (C-D) Representative images of 3×FLAG-HRJDa/b localization in wing discs and adult midguts. FLAG-tagged HRJDs were induced using *WP-Gal4* (C) or *esg-Gal4* (D). Scale bars: 10 µm.

**Figure S2. Continuous expression of HRJDs throughout development hampers intestinal regeneration**

(A-C) Organismal survival upon paraquat (A), DSS (B), and sucrose (C) feeding. Drug feeding started at Day 7 after eclosion. HRJD induction using *esg-Gal4* throughout development significantly decreased survival upon paraquat/DSS feeding.

(D) Expression of HRJDa using *esg-Gal4* decreased the number of PH3 positive cells in the posterior midguts, while HRJDb did not significantly change PH3 positive cell number.

N.S., not significant: P>0.05, **P≤0.01, ***P≤0.001, one-way ANOVAs with post hoc Tukey test. *n* indicates the number of flies (A-C) and guts (D).

**Figure S3. Organismal lifespan of flies expressing HRJDs in ECs and ISC/EBs**

(A-F) Survival curve for organismal lifespan by sex. HRJDs are induced by *NP1-Gal4*

(A and B), *NP1^ts^-Gal4* (C and D), and *esg-Gal4* (E and F).

N.S., not significant: P>0.05, ***P≤0.001, Log-rank tests. *n* indicates the number of flies.

**Figure S4. HRJD induction in adult ISCs/EBs promotes turnover of midgut epithelial cells.**

(A) Representative images of esgFLPout aged midguts in which *esg*-lineage clones express HRJDs together with GFP. Clones were induced 50 day after eclosion and traced for 10 days (final 60-day old adults). Ssk was immunostained to compare the septate junction of HRJD-expressing cells with that of wildtype cells, which is quantified in (B). The magnified images of areas enclosed by squares are shown as images 1-5. Arrows indicate GFP^+^ polyploid adjacent cells in the control midgut.

(B) HRJD expression improved Ssk accumulation at septate junction not only in HRJD-expressing cells (GFP^+^/GFP^+^), but also in wildtype cells (GFP^−^/GFP^−^).

(C) HRJD expression increased the ratio of GFP^+^ cells and total cells, suggesting that old ECs are more rapidly replaced by newly formed ECs.

(D) HRJD expression increased the ratio of GFP^+^ polyploid cells and GFP^+^ diploid cells, suggesting that HRJDs increases the speed of differentiation to ECs.

(E) Quantification of PH3 positive cell number in Day 10 young posterior midguts. HRJD induction with *esg-Gal4, tub-Gal80ts* driver did not change the quantity of mitotic cells. (F-G) HRJD induction increased the ratio of cDCP1^+^ cells in aged midguts (F), which is quantified in (G).

(H) RT-qPCR of signaling ligands established as mitotic activators in the adult midgut.

N.S., not significant: P>0.05, *P≤0.05, **P≤0.01, ***P≤0.001, one-way ANOVAs with post hoc Tukey test. *n* indicates the number of cells (B), midguts (C, D, E and G), and biological replicates (H). Scale bar: 50 µm or 25 µm (magnified images 1-5).

**Table S1.**
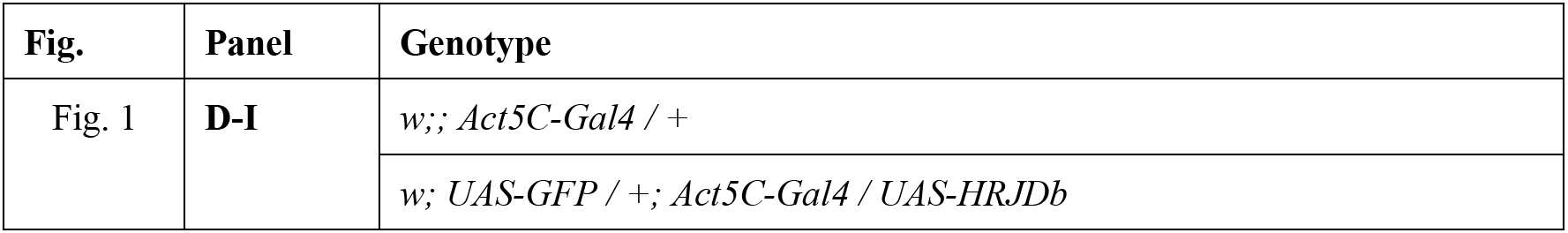
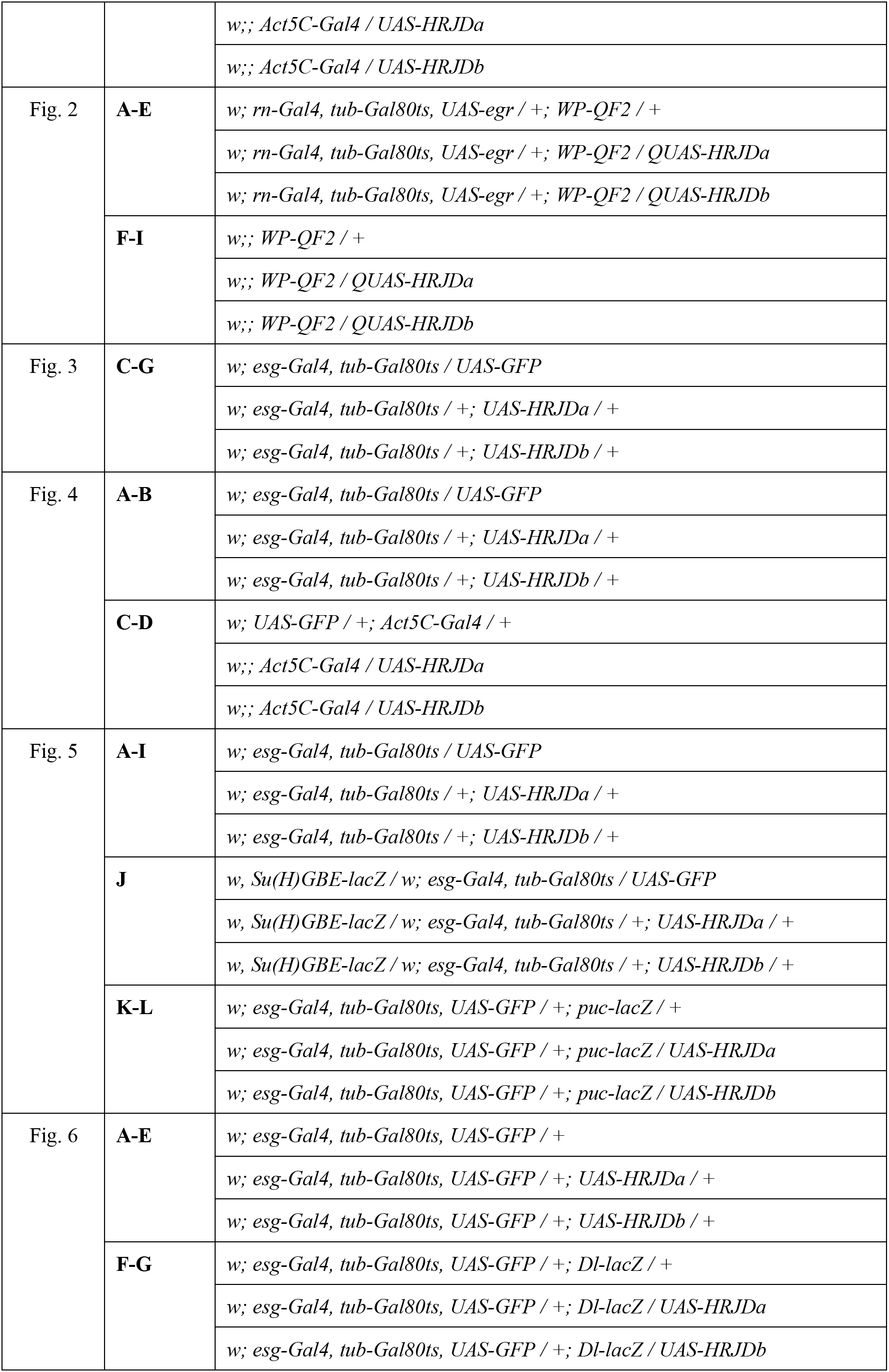
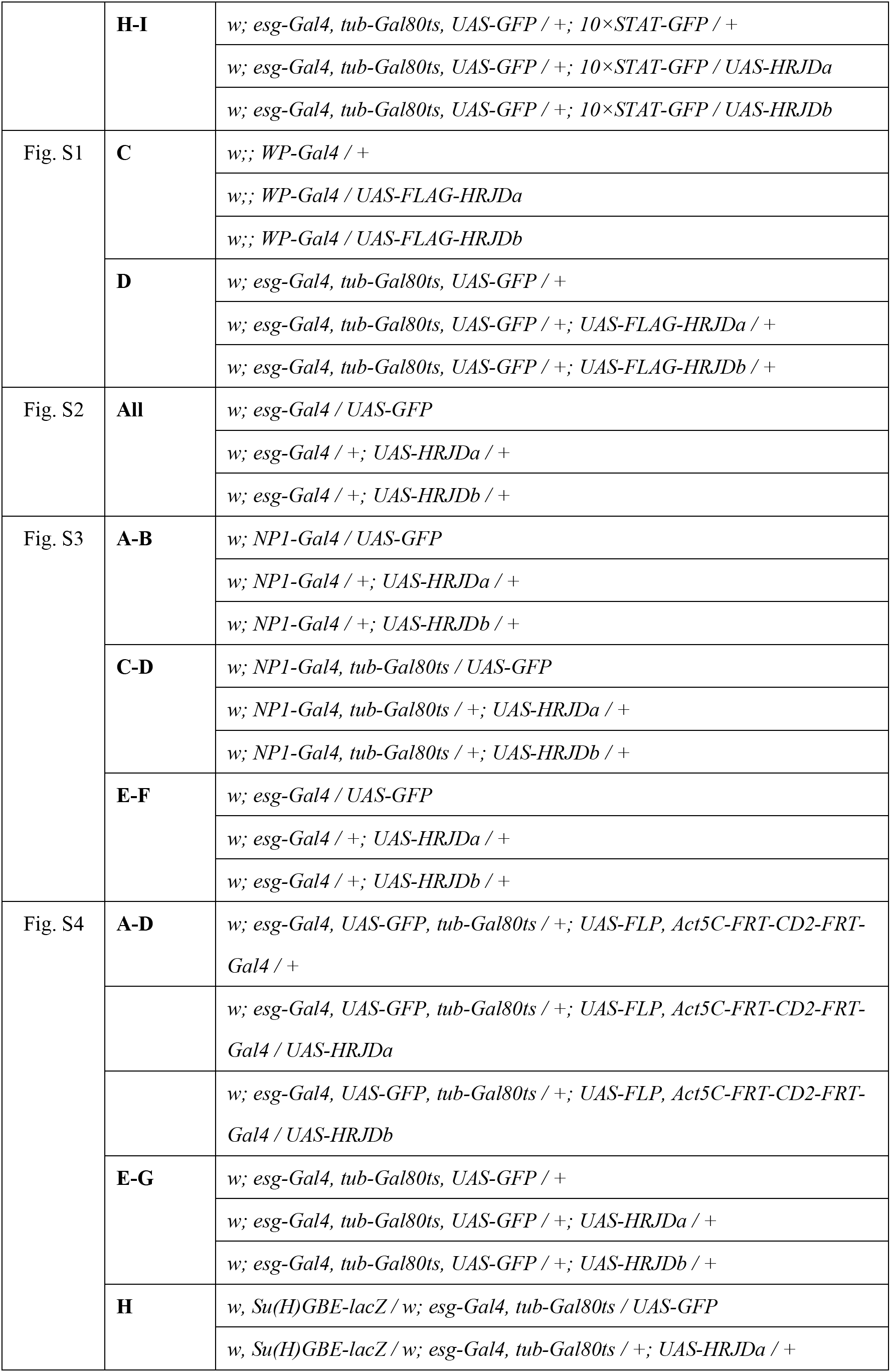
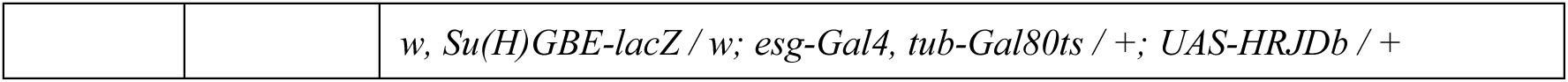
Detailed genotypes in each experiment.

**Table S2. Primer sequences**

**Supplementary texts S1. Sequences for pUCIDT-HRJDa and pUCIDT-HRJDb**

## Notes

### Competing Interest Statement

The authors have declared no competing interest.

### Summary of Updates

The mechanism of HRJD-mediated ISC rejuvenation was proposed; HRJD in ISCs promotes EC differentiation and cellular turnover; Figure 6 revised; Supplemental Figure S4 added.

