## Supplementary figures and images for "Highly regenerative species-specific genes improve age-associated features in the adult *Drosophila* midgut"

### Supplementary information

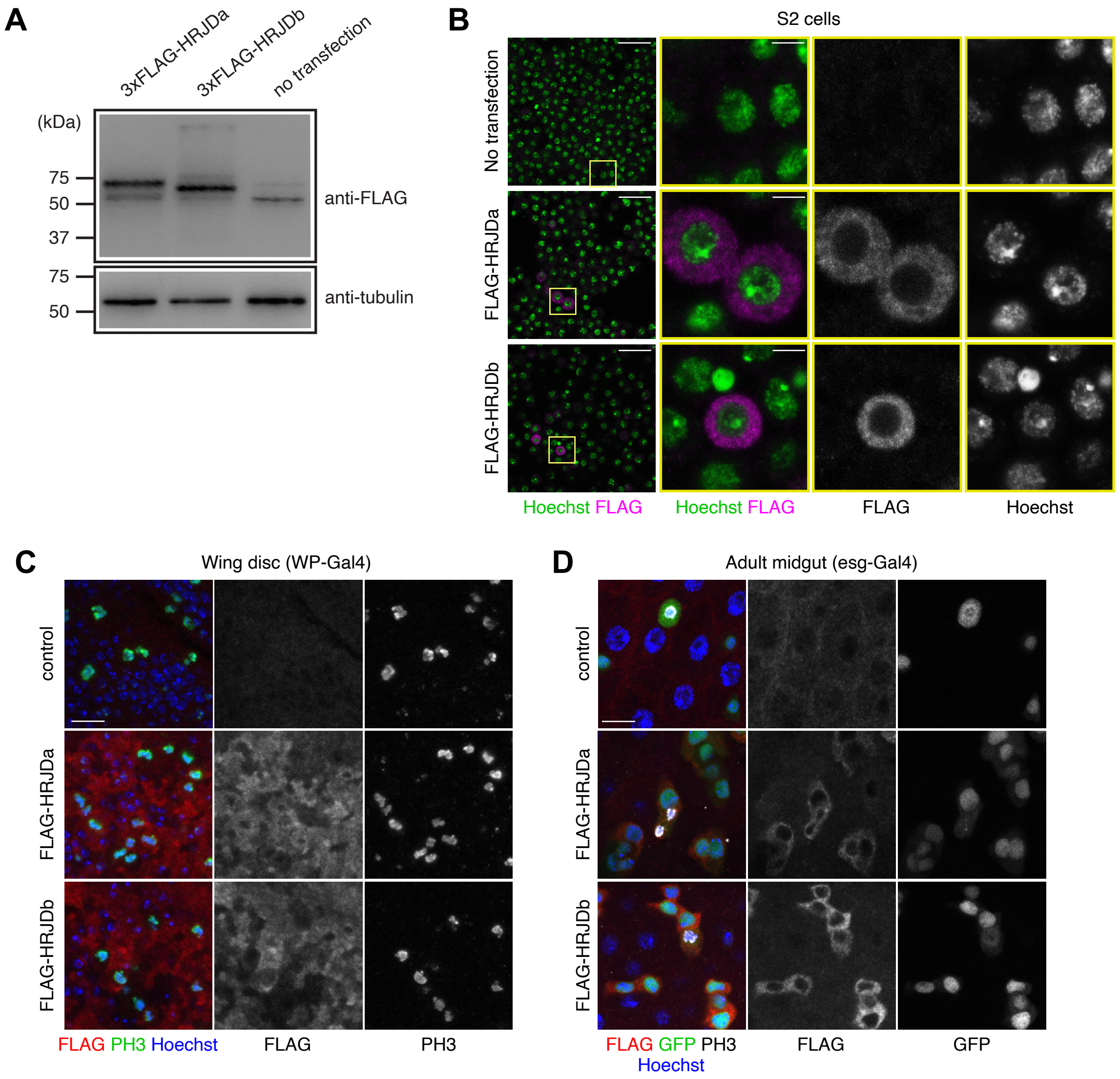

Nagai et al.

Figure S2

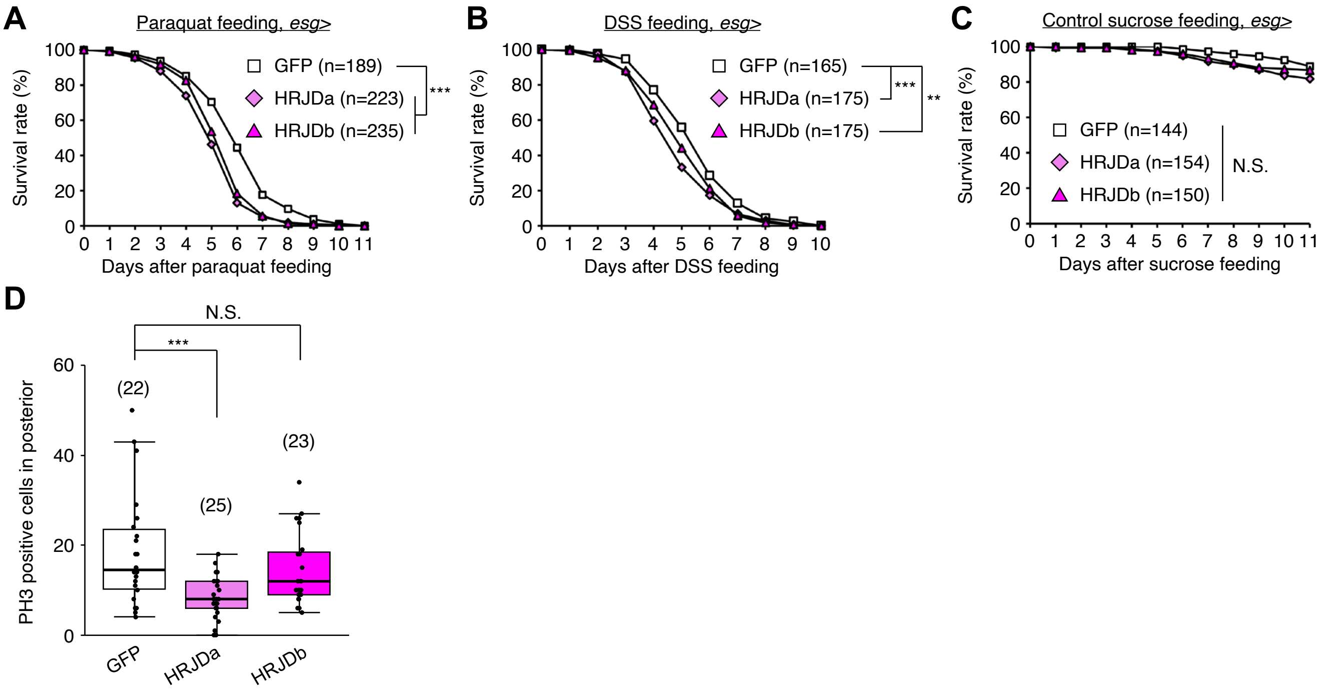

**Nagai et al.** **Figure S3**

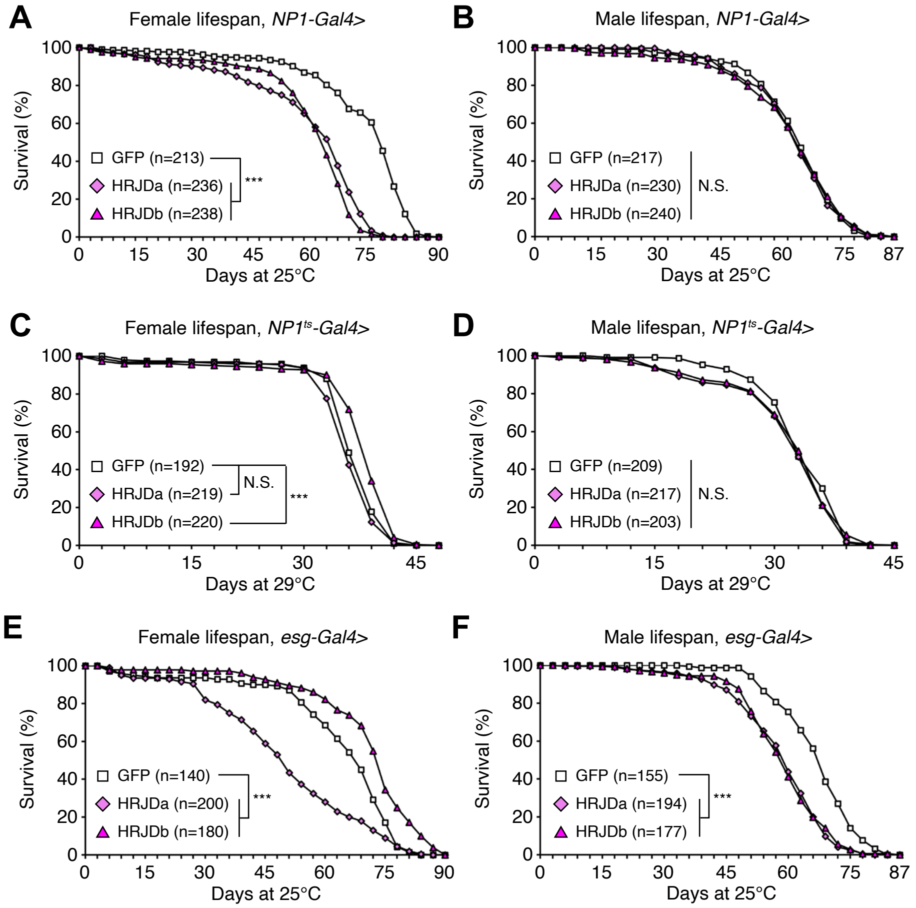

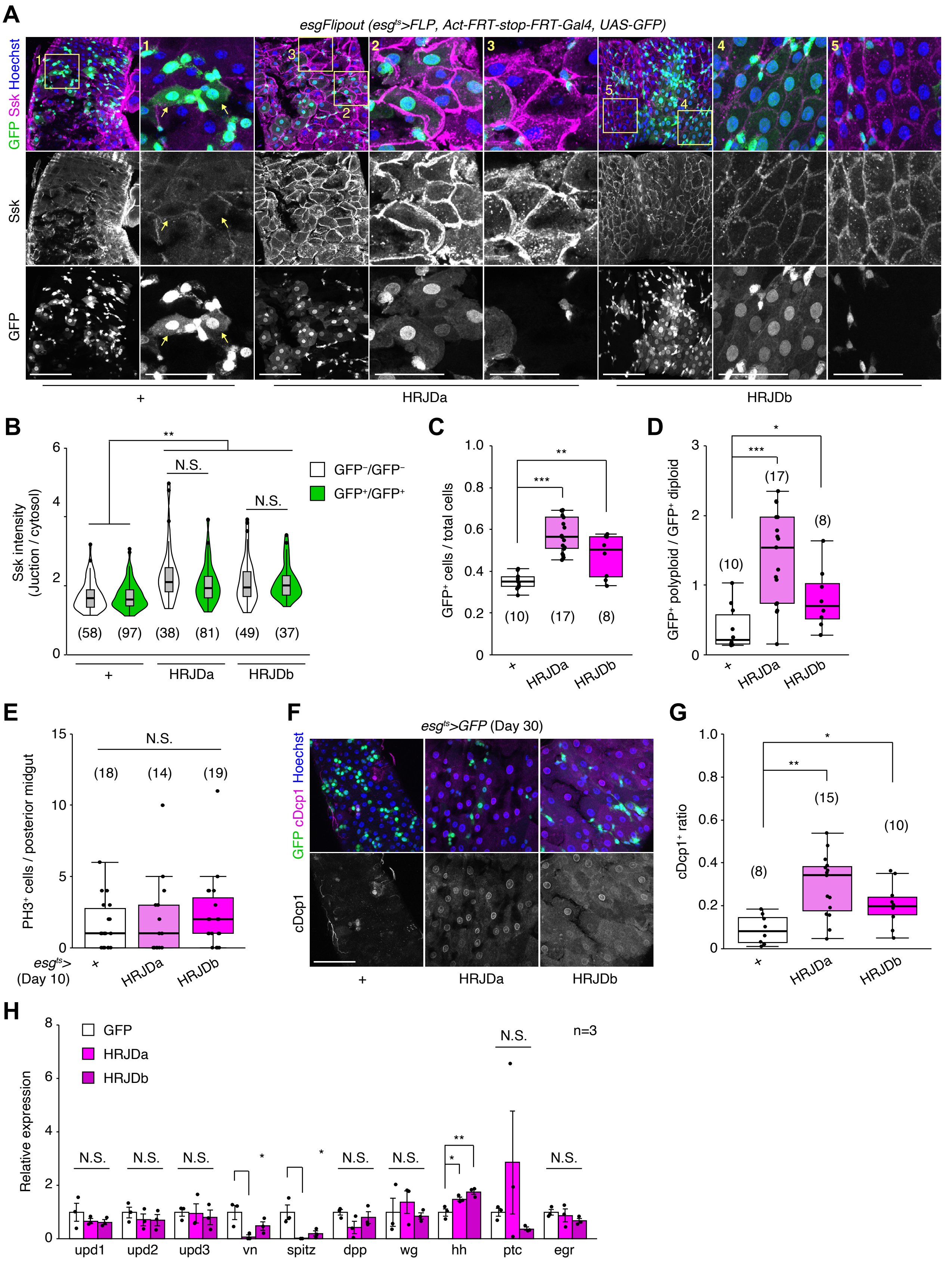
